# Primitive and modern swimmers solve the challenges of turning similarly to achieve high maneuverability

**DOI:** 10.1101/706762

**Authors:** J. O. Dabiri, S. P. Colin, B. J. Gemmell, K. N. Lucas, Megan C. Leftwich, John H. Costello

## Abstract

Animal swimmers alter trajectories – or turn - for a variety of critical life functions such as feeding, mating and avoiding predation. Yet turning represents a fundamental dilemma based in rotational dynamics: the torque powering a turn is favored by an expanded body configuration, yet minimizing the resistance to a turn (the moment of inertia) is favored by a contracted body configuration. How do animals balance these opposing demands to achieve high maneuverability? By noninvasively measuring fluid and body motions, we found that both jellyfish (*Aurelia aurita*) and fish (*Danio rerio*) initially maximized torque using previously undescribed, rapid body movements. Both species then minimized resistance to turning by bending their bodies to reduce their moment of inertia. Use of this sequential solution by such distantly related animals as an invertebrate and a vertebrate suggests strong selection for these turning dynamics that may extend to other swimmers and inform future vehicle designs.

## Introduction

The study of aquatic locomotion has primarily examined the dynamics and energetics of linear, unidirectional swimming (*1–14*). However, this focus on parameters governing linear, unidirectional swimming obscures a more subtle reality that animals swimming in nature rarely follow simple linear trajectories. Instead, natural animal paths typically involve frequent changes in direction that are mediated by turning maneuvers. The widespread importance of these turning events is evident in the range of models describing circuitous natural animal pathways (*15–17*). These studies make clear that, from microscopic plankton (*18*) to humpback whales (*19*), swimming animals exhibit predominantly shifting pathways with frequent turns which alter their trajectories. The complexity of animal swimming under natural conditions demonstrates that rotational motions as well as linear translation are fundamental components of animal swimming.

Although central to natural patterns of animal movement, the process of turning entails a fundamental and unaddressed mechanical challenge for animal swimmers. The challenge is one of rotational physics, namely, the same body configurations that maximize the torque for turning also maximize that body’s resistance to turning. Specifically, torque increases in direct proportion with the distance from the center of rotation at which locomotive force is applied. For swimming animals, this lever arm is primarily set by the distance of the swimming appendages from the body center of mass. The resistance to turning, which is characterized by the body moment of inertia, increases even more significantly with an increasing lever arm, exhibiting a quadratic dependence (see mathematical treatment in Method). The body moment of inertia has been demonstrated to be a key factor affecting both animal (*20*, *21*) and human (*22*, *23*) angular velocities during turns in air and water. It remains unclear how animal swimmers resolve the conflicting demands of high torque production (i.e. expanded body configuration) with those of low moment of inertia (i.e. contracted body configuration) to achieve high turning performance. Knowledge of such solutions is important for understanding maneuverability by swimming animals, and successful strategies may potentially inspire high maneuverability in engineered vehicles.

To evaluate this question broadly, we chose two model species with extremely divergent body types, neural organization, and phylogenetic relatedness. The jellyfish *Aurelia aurita* is a member of the oldest animal group to use muscle fiber-driven swimming and one of the most energetically efficient metazoan swimmers. *A. aurita* is characterized by a radially symmetrical body plan and a very restricted array of cell types with comparatively simple neurological organization. Scyphomedusae such as *A. aurita* lack true muscle cells found in other phyla and, instead, rely upon only a thin sheet of striated muscle fibers located within epithelial cells for body movements (*24*). In contrast, the zebrafish *Danio rerio* represents the evolution of a bilaterally symmetrical body plan with comparatively complex skeletal structure and neuromuscular organization characteristic of modern fish species (*25*).

Experiments were conducted in transparent laboratory vessels filled with seawater. Animals were studied swimming individually in the tank in the absence of ambient currents. Measurements of the flow induced by the animals were non-invasively collected using particle image velocimetry (PIV). A horizontal laser sheet illuminated 10 micron, neutrally-buoyant glass beads suspended in the water. A high-speed camera simultaneously recorded the animal swimming kinematics and the motion of the glass beads. Only sequences in which the animal’s symmetry plane coincided with the plane of the laser sheet were used for analysis. The velocity fields measured using PIV were subsequently input to custom algorithms to compute the corresponding pressure fields and forces surrounding the animal (*26, 27*), see Methods for further details.

## Results

Jellyfish (Fig 1A-L)-and zebrafish (Fig. 1M-X) both exhibited frequent bouts of turning, during which flow measurements revealed pronounced changes in the pressure fields in the water adjacent to the animal (Figs. 1F and 1R, for jellyfish and fish, respectively). These significant pressure fields preceded the more pronounced body motions that occurred during the subsequent turn that changed the animal swimming direction (Figs. 1C and 1O, respectively). Examination of the body shape during the period of transient pressure buildup led to the discovery of a small, rapid shift in the curvature of the animal body immediately preceding the turn for both the jellyfish (1.5 ± 1.0 percent change in curvature, n = 10) and the zebrafish (0.8 ± 0.2 percent, n = 10). Although the amplitude of this initial body bend was small, it occurred over a sufficiently short period of time - a few milliseconds - that the corresponding acceleration of the body was large relative to accelerations during unidirectional swimming. The measured peak accelerations preceding the turn were over 1 m s^−2^. This motion was transmitted to the adjacent water via a process known as the acceleration reaction or added-mass effect (*28*) (Fig. 2).

**Figure 1.**
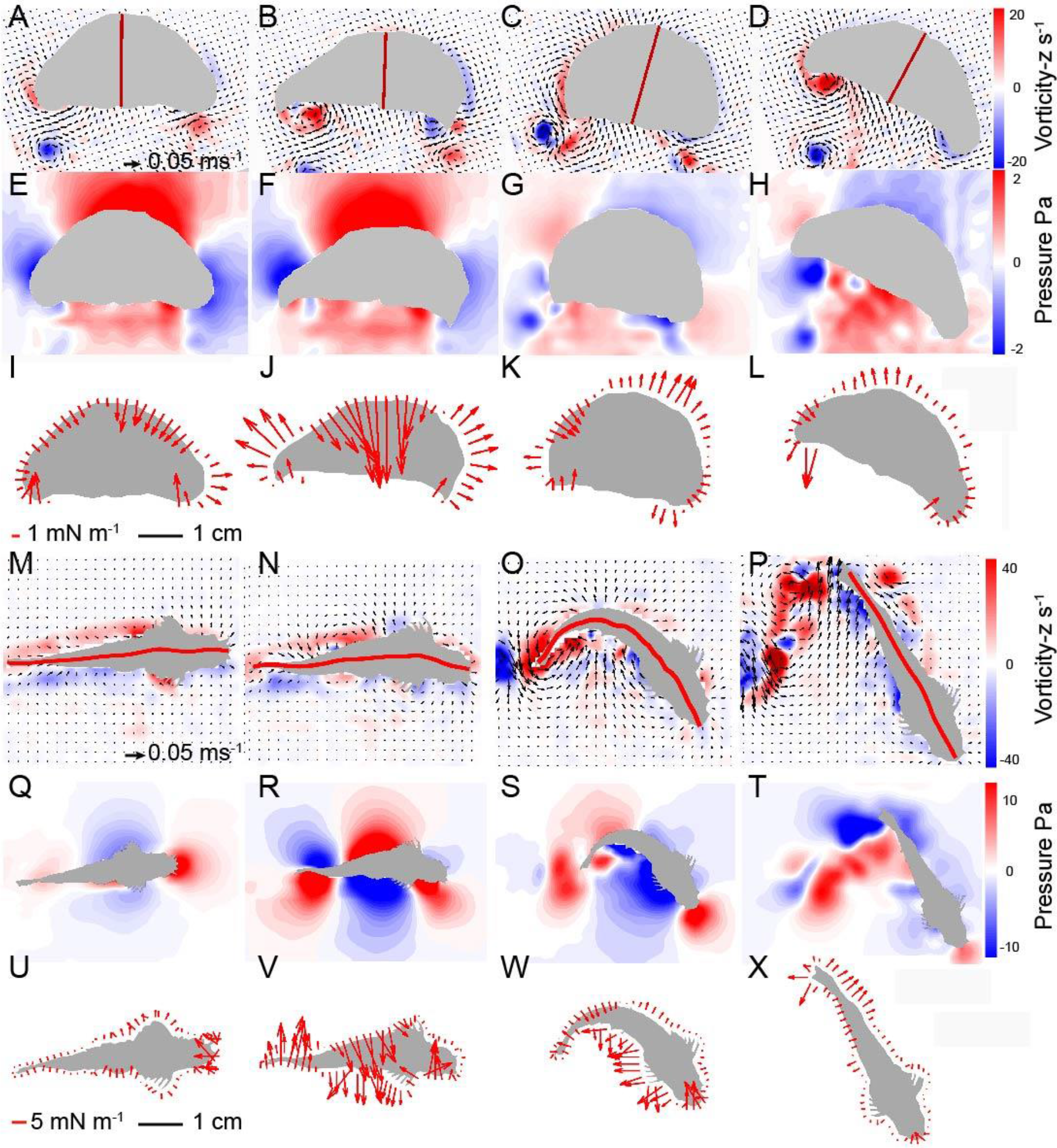
Fluid interactions and forces during turning by jellyfish and fish. Sequential panels describe the turning kinematics and fluid pressure for representative medusa (*Aurelia aurita*, 30° rotation, profiled in Fig. S1d) and zebrafish (*Danio rerio*, 62° rotation, profiled in Fig. S2c) turns. The red line shows the midline of the medusa (**A-D**) and the fish (**M-P**) throughout the turn, along with PIV vector and vorticity fields. Pressure fields around the medusa (**E-H**) and the fish (**Q-T**) demonstrate that both animals generate large, asymmetric pressure gradients around their bodies (panels **F** and **R**, respectively) before major body orientation shifts (illustrated by the midline position). Force vectors due to local fluid pressure at the medusa (**I-L**) and zebrafish (**U-X**) body surface indicated in red arrows. Note the location and magnitude of force vectors along the bodies of jellyfish (**J**) and fish (**V**) that generate high torque while the animal’s body is extended and before major body-bending occurs. Supplementary movies illustrate examples of velocity fields, pressure and force for medusae (Movies S1-S3) and zebrafish (Movies S4-S6).

**Figure 2.**
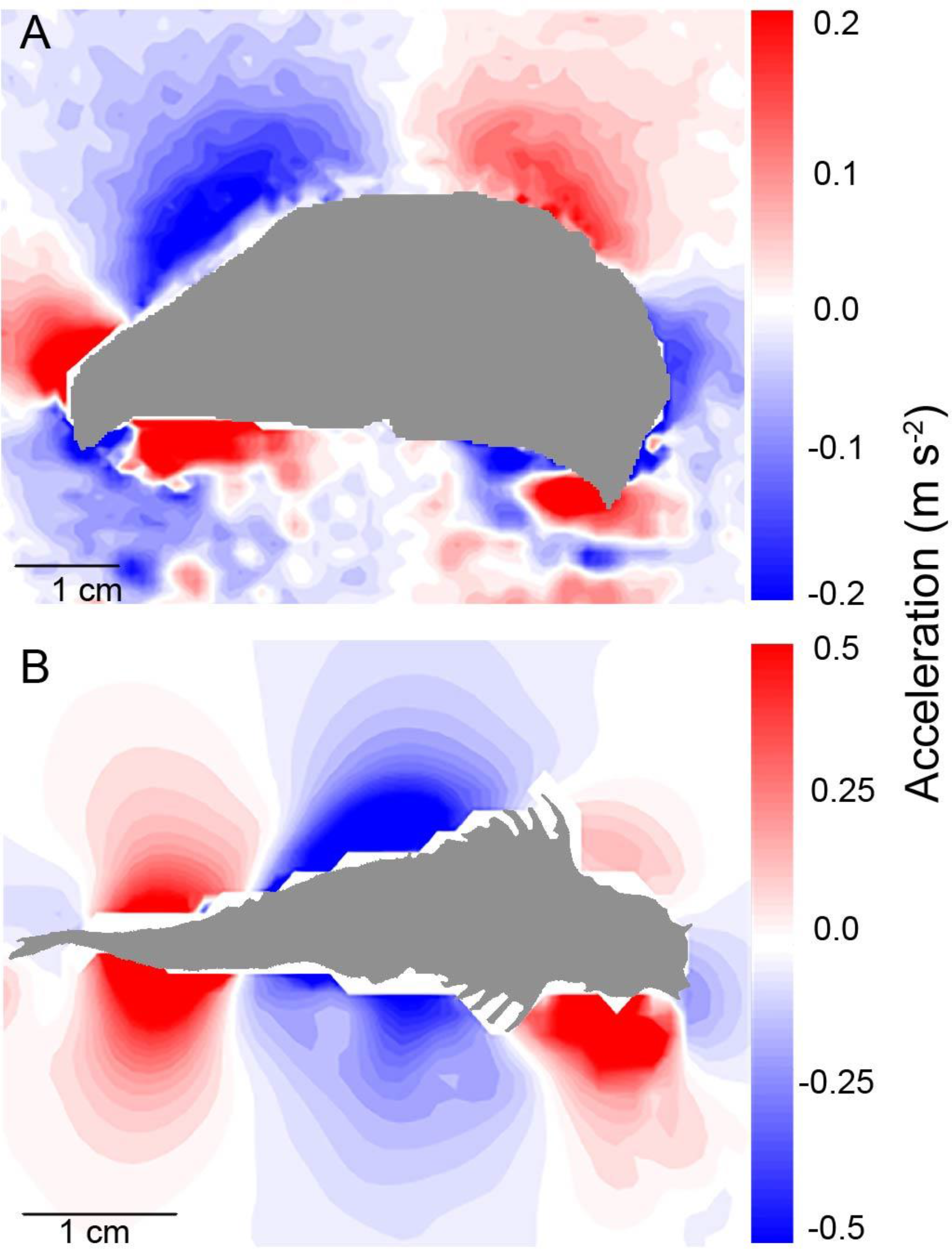
Rapid fluid accelerations during turn initiation give rise to high torque forces along the bodies of jellyfish and fish. Fluid acceleration (positive values correspond to vertical motion toward bottom of page) along animal bodies during turn initiation by medusa (**A**, *Aurelia aurita*) and zebrafish (**B**, *Danio rerio*). Fluid accelerations in both panels are for the same turning sequences as depicted in Figure 1, so that the acceleration field in panel **A** corresponds to the high pressure state of Fig. 1**F**, while panel **B** corresponds to that of Fig. 1**R**.

Because the water is effectively incompressible, the fluid in contact with the body responded to the high local body acceleration by an increase in the local fluid pressure where the body was advancing (pushing the water), and a decrease in the local pressure where the body surface retreated from the local water (pulling the water with it). When integrated over the full animal body, the pressure field created by the small, asymmetric body bending leads to a large net torque capable of turning the organism toward a new heading. The more pronounced body bending that occusr after the generation of this pressure field does not contribute greatly to torque generation, but does reduce the moment of inertia of the body (Fig. 3; see also Figs. S1 and S2). Therefore, the body kinematics that follow peak pressure generation enhance the effect of the generated torque by amplifying the resulting angular acceleration so that the body axis rotates rapidly through a turn. This sequence of body kinematics that initially maximizes torque forces and subsequently minimizes the moment of inertia resolves the fundamental competition between these two components of rotational motion during turns. Although the maximum torque generation and minimum moment of inertia do not occur simultaneously (Fig. 4), the inertia of the fluid and of the animal body allows the initial pressure transient to affect subsequent turning dynamics.

**Figure 3.**
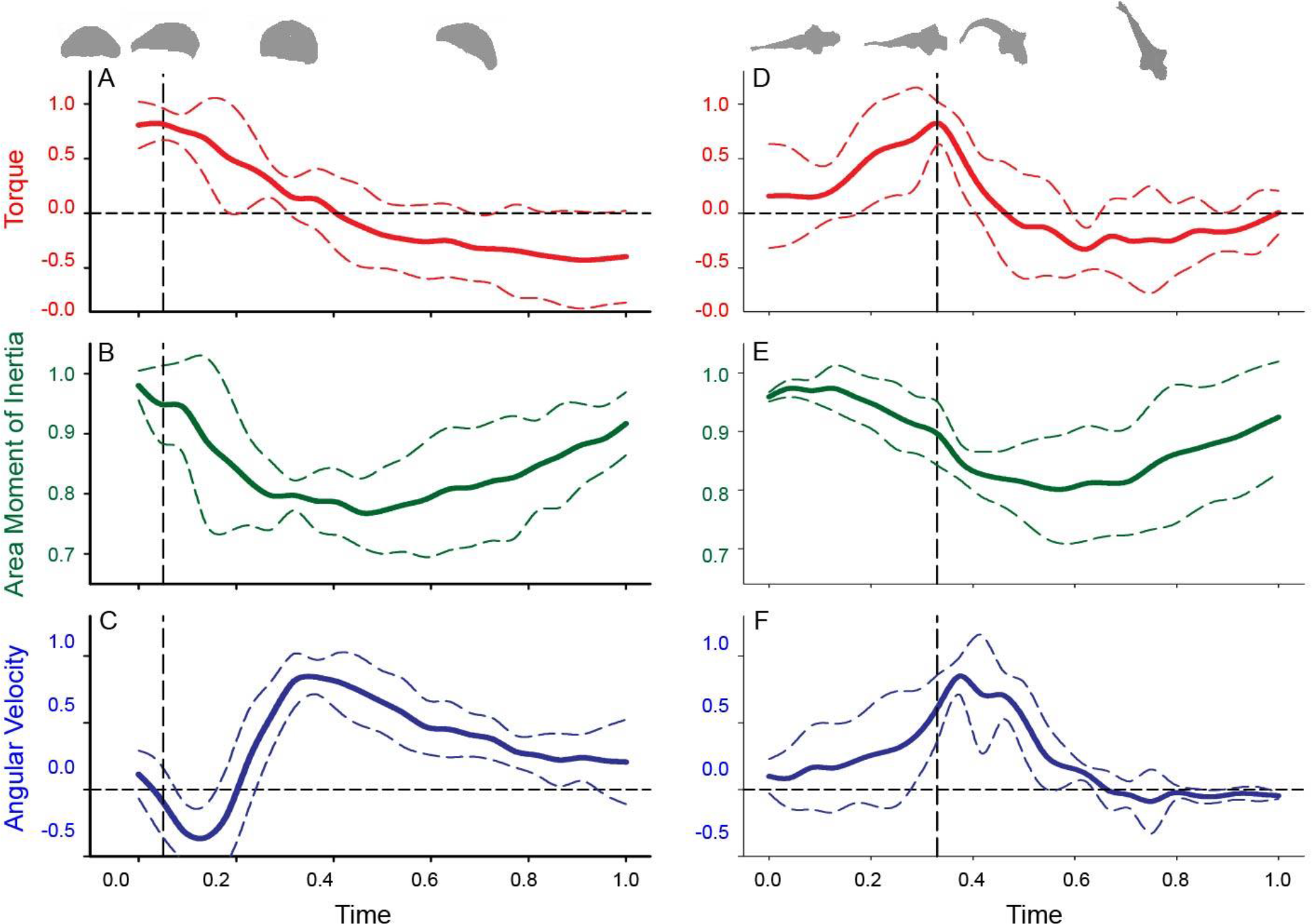
Normalized data for comparison of turning variables between jellyfish and fish. Patterns represent data for replicate individuals during variable turn excursions (medusa *Aurelia aurita*, panels **A-C**, n = 6; bell diameters 1.8-5.4 cm, range in turn angles 13-53°; zebrafish *Danio rerio*, panels **D-F**, n = 5, fish lengths 3.6-4.7 cm, range in turn angles 17-95°). Data for each replicate turn was divided into a uniform number of sample intervals and each variable (time, area moment of inertia, angular velocity and torque) was normalized by the highest value of each replicate sequence so that all variables could be expressed in dimensionless form with a maximum value of 1. Vertical dashed black line represents the time of peak torque production. Solid colored curves represent the mean value and dashed lines represent one standard deviation above or below the mean for each sample interval. Note that peak values do not always reach 1 because they are averages of all the turns and not all the peak values occurred in the same time interval for every turn. The original, non-normalized data for each individual replicate are displayed in Fig. S1 (jellyfish) and Fig. S2 (zebrafish).

**Figure 4.**
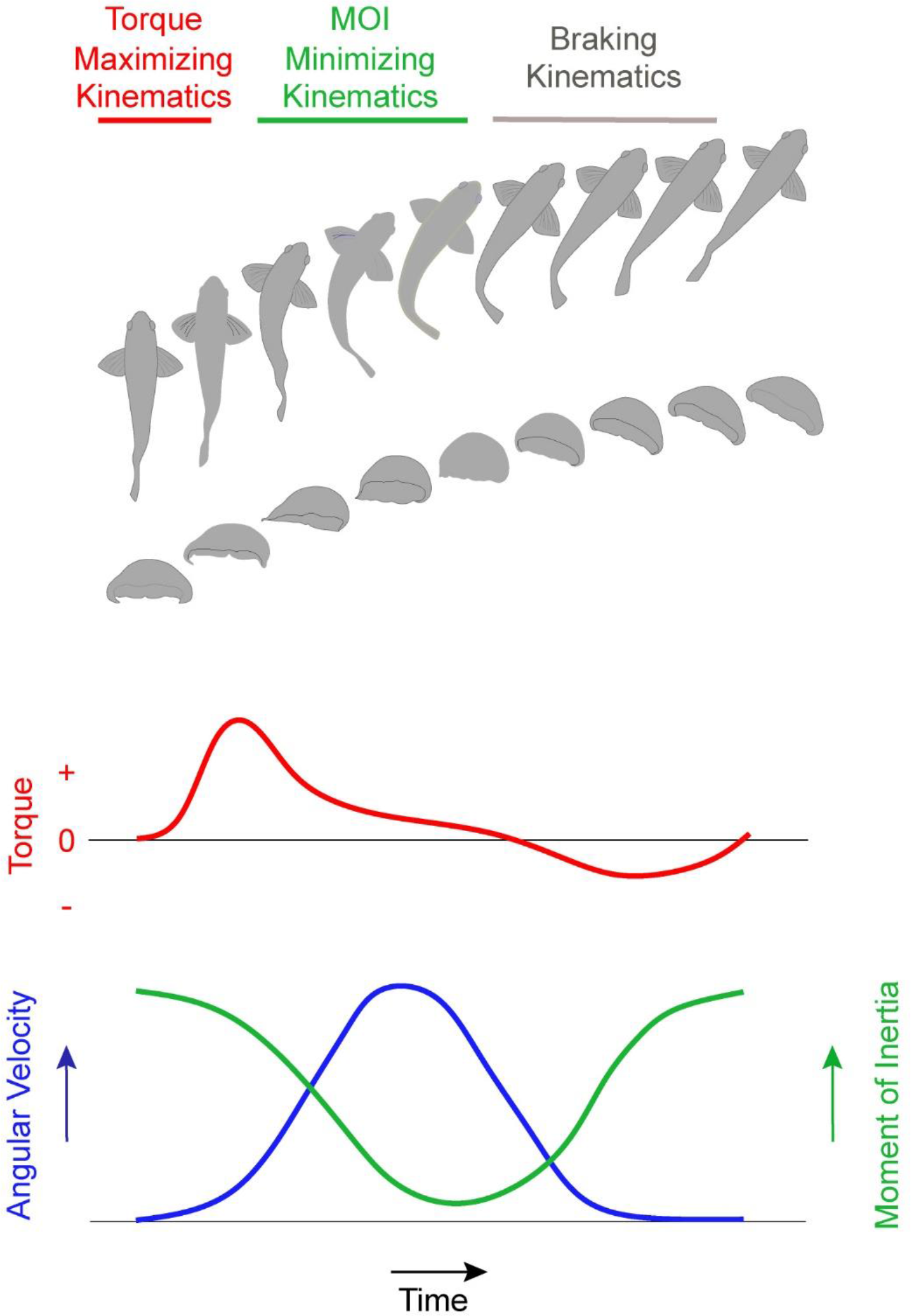
Conceptual summary of turning dynamics by the jellyfish (*Aurelia aurita*) and the zebrafish (*Danio rerio*). Arrows for each axis represent increasing magnitude for that variable. A turn is initiated by a subtle body bend which builds torque before the animal turns (changes heading). After peak torque production, the animal bends its body more radically to minimize its moment of inertia (MOI). This decreases the body’s resistance to rotational motion while increasing angular velocity and turning the animal. The turning sequence ends as negative torque brakes the turning rotation when the body returns to its extended configuration with high moment of inertia and low angular velocity.

## Discussion

We observed strikingly similar turning dynamics for both the jellyfish and the zebrafish, despite their substantially different body organization and swimming kinematics (Figs. 1–3). The dynamical importance of the observed pressure fields for both the jellyfish and zebrafish was confirmed by computing the net torque (per meter depth) and area moment of inertia of the body. For turns of varying net change in heading, the initial pressure pattern created by the animals was nearly constant. The ultimate magnitude of each turning maneuver was instead modulated by changes in body shape that tuned the moment of inertia and thereby controlled the angular acceleration of the body. In all cases, the relationships between pressure measurements and turning kinematics followed a similar sequential pattern (Fig. 3 for average patterns, details for all replicates in Figs. S1 and S2).).

An essential feature of animal turning by the mechanisms described here is the flexibility of the body, which enables the animal to dynamically redistribute its mass to manipulate the lever arm of the propulsive surfaces used to initiate the turn (e.g. the bell margin of the jellyfish and the caudal fin of the zebrafish) and the body moment of inertia (Fig. 4). For animal swimmers with flexibility and size scales favoring this process, the performance advantages of this turning strategy may select for very similar turning kinematics despite the vastly different animal forms studied here.

While the present results motivate further study of turning in other swimming animals whose locomotion lies between jellyfish and zebrafish, we anticipate that extension of these findings will depend upon scaling factors that influence the size range over which this approach is effective. In the regime of swimming at low Reynolds numbers (Re = *ULv*^−1^, where *U* and *L* are the nominal animal swimming speed and size, respectively, and *v* is the kinematic viscosity of the water), angular momentum generated during periods of maximum torque would experience rapid viscous dissipation, leaving little remaining angular momentum to complete the turn during the subsequent period of major body bending. For large animals with body lengths on the order of tens of meters, power requirements for rapid body bending may exceed the available muscle capacity. In geometrically similar animals, angular acceleration scales to the −2/3 power of body mass (*29*), making it more difficult for large animals to generate the initial pressure transient or to alter their moment of inertia through body rearrangement to increase their angular velocity. Hence, very large swimmers such as whales may not bend as readily as smaller animal swimmers such as zebrafish. However, the majority of animal swimmers exist within the millimeter to meter size range (*30, 31*), in which a time-varying lever arm enabled by body bending would provide favorable performance advantages relative to rigid body turning mechanics.

These observations of a large dynamical impact from small kinematic shifts compel further study of the neuromuscular control of aquatic locomotion and engineered systems that aim to be inspired by animal swimming. In particular, while nature has not converged upon unidirectional locomotion that leverages similar kinematic subtleties as in turning (i.e. steady, straight swimming does not exhibit the small body motions observed here), it might be feasible to achieve net propulsion using such an approach in a robotic system. The pronounced pressure fields observed presently in the jellyfish and zebrafish are incompatible with unidirectional translation, as they achieve high net torque but low net force due to the balance of high and low pressure on either side of the animal. However, it is conceivable that modified kinematics could result in net propulsive force. Perhaps the greatest import of these turning mechanics lie with potential vehicle maneuvering performance. Animals swimmers are characterized by rotational velocities that are substantially higher than rigid human designed vehicles and consequently, have much higher turning rates(*32*). The underlying mechanics of turning by animal swimmers may provide a useful blueprint for more maneuverable vehicle designs employing soft materials (*33*).

More broadly, an appreciation of the important role of turning maneuvers in the success of aquatic locomotion can reshape ongoing and future efforts to understand the role of physical forces in the evolution and ecology of both primitive and modern swimmers. The methods employed here to study freely swimming organisms and to quantify their dynamics in terms of pressure field manipulations provide a powerful tool to enable new insights into aquatic locomotion. This solution arrived at by such different animal lineages allows them to initially maximize torque production before major body curvature change and subsequently minimize the moment of inertia by bending. The effectiveness of this solution for rotational motion coupled with the pervasive demands of turning provides insight into the near universal capability of swimmers to re-arrange their mass by flexible bending. The subtleties of this pattern have been obscured by the historical focus on parameters governing linear, unidirectional swimming but may prove more important for explaining the evolution of efficient aquatic locomotion employed by animals in their natural environments.

## Materials and Methods

### Animals and imaging

The zebrafish (*Danio rerio*) used in this study were adults acquired from the Zebrafish Facility at the Marine Biological Laboratory (MBL). All procedures were in accordance with standards set by the National Institutes of Health and approved by the Institutional Animal Care and Use Committee at the MBL. Zebrafish were maintained at room temperature (23-25°C) in 37 l aquaria until imaged while swimming. Swimming and turning behaviors (n = 5) were recorded as individual fish swam along the center of an acrylic raceway tank (1.5 × 0.5 m). *Aurelia aurita* medusae were obtained from the New England Aquarium and maintained at 25°C in 20 l aquaria. Single, representative animals (n = 6) were recorded while freely swimming in a 0.3 × 0.1 × 0.25 m glass vessel, using methods reported previously^11^.

### Particle image velocimetry (PIV)

We used high-speed digital particle image velocimetry (PIV) to obtain resulting flow fields around the fish and medusae. Recordings were acquired by a high-speed digital video camera (Fastcam 1024 PCI; Photron) at 1000 frames per second and at a spatial resolution of 1024 × 1024 pixels with a scale factor of 0.178 mm per pixel. Seeding particles (10 μm hollow glass beads; Potters Industries) were laser-sheet illuminated for PIV measurements. Medusae were illuminated with a laser sheet (680 nm, 2W continuous wave; LaVision) oriented perpendicular to the camera’s optical axis to provide a distinctive body outline for image analysis and to ensure the animal remained in-plane, which ensures accuracy of 2D estimates of position and velocity. The transparent bodies of medusae allowed a single laser light sheet to illuminate fluid surrounding the entire body. Fish were not transparent and so were illuminated by two laser sheets (532 nm, 600 mW continuous wave, Laserglow Inc.) mounted in the same plane on opposite sides of the tank to eliminate shadows on either side of the body as each animal swam within the field of view^25^.

Fluid velocity vectors for both fish and medusae were determined from sequential images using a cross-correlation algorithm (LaVision software). Image pairs were analyzed with shifting overlapping interrogation windows of a decreasing size of 32×32 pixels to 16×16 pixels. Masking of the body of the fish before image interrogation confirmed the absence of surface artifacts in the PIV measurements.

### Pressure and torque measurement

Velocity fields collected via PIV were input to a custom program in MATLAB that computed the corresponding pressure fields. The algorithm integrates the Navier-Stokes equations along eight paths emanating from each point in the field of view and terminating at the boundaries of the field of view. The pressure at each point is determined by computing the median pressure from the eight integration results. Bodies of the fish and medusae were masked prior to computation to prevent surface artefacts in the pressure and torque results. Masks were generated using a custom MATLAB (Mathworks, Inc.) program that automatically identified the boundary of the animal body based on image contrast at the interface between the animal body and the surrounding fluid, and body outlines were smoothed prior to later analyses. These methods have been previously validated against experimental and computational data, including numerical simulations of anguilliform swimming (*27*) and direct force and torque measurements of a flapping foil (*26*). The MATLAB code is available for free download at http://dabirilab.com/software.

The fluid force normal to the body surface due to the local fluid pressure was determined by integrating the calculated pressure along the corresponding surfaces of the body (*26*). Validations against measurements made on physical models show that these calculation techniques based on 2D PIV images are robust to a small degree of out-of-plane flow such as that induced by a fish’s slight rolling motions during turns, so long as the fish remains centered in the imaging plane (*26*). The body outline of each animal was divided into segments of equal length (zebrafish: 84 segments, medusa: 70-85 segments) for spatial integration. For medusae, the laser light sheet passed through the central axis of the body, but surface segments defined along the central bell margin run outside of the laser light sheet. These bell components outside the laser sheet can interfere with images of the bell cavity. Although these surface segments were required to mask the animal body, they did not represent surfaces visible within the PIV laser light sheet, and so forces and torques calculated on these central bell segments were not included in later analyses.

Because the surface geometry was specified in a single plane, the force calculations were evaluated per unit depth (i.e. giving units of Newtons per meter of depth perpendicular to the measurement plane). The corresponding torque was calculated as the vector product of the moment arm from each location on the body surface to the center of mass, and the local force due to pressure at the same location on the body surface. The resulting torque calculations have units of Newton-meters per meter, corresponding to the aforementioned planar measurements. MATLAB codes for force and torque calculations similar to those conducted presently as well as the segment-making methods are available on Github (https://github.com/kelseynlucas) and have been validated in earlier work (*26*).

### Turning equations of motion

The mass moment of inertia of a body is a measure of how its mass is distributed relative to a reference axis, often taken as the geometric centroid. It is given by

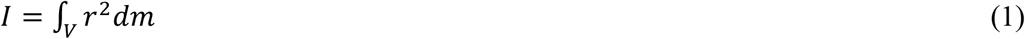

where *V* is the region occupied by the body mass, and *r* is the distance of each infinitesimal portion of body mass from the reference axis. In the present case, this mass moment of inertia was approximated using the area moment of inertia, which is a measure of how the body area in a cross section is distributed relative to the reference axis:

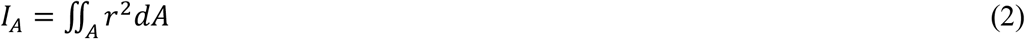

where *A* is the region occupied by a two-dimensional cross-section of the body. The cross-section in the present measurements was the body symmetry plane illuminated by the laser sheet during PIV measurements. The area moment of inertia (henceforth called the moment of inertia for brevity) was calculated using a custom program in MATLAB as described in the following section.

The torque exerted on a body is related to changes to both its angular motion and its moment of inertia by the following relation:

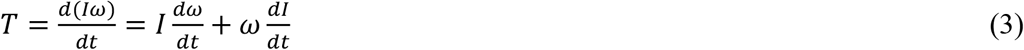

where *ω* is the angular velocity of the body. The first term of the summation incorporates the rate of change of angular velocity, i.e. the angular acceleration. The second term depends on the change in the moment of inertia, i.e. changes in body shape or mass. As illustrated in comparison of Fig. 3a-c with Fig. 3d-f, different temporal trends of *I*, *ω*, and their time-derivatives can be consistent with the measured net torque via application of equation (3).

### Moment of inertia and angular velocity measurements

Calculations of the moment of inertia for turning sequences used the same smoothed animal body outlines automatically detected for pressure and torque calculation. A separate custom MATLAB algorithm subsequently calculated the moment of area for each image. Their bodies were partitioned as for the force and torque measurements above, with each of segment of area *a*_*i*_ having a centroid located at distance *r*_*i*_ from the whole body centroid. The area moment of inertia for each frame *p* was then calculated as:

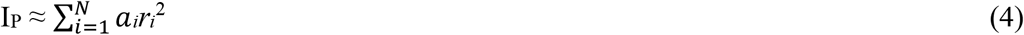

where the summation was taken over the *N* body segments. Angular velocities of zebrafish during turns used local body surface position changes to calculate the angle of the line segment connecting the anterior head region with that of the body centroid. The rate of change of that angle in a lab-fixed frame determined the fish angular velocity. The hemi-ellipsoidal shape of medusae and shifts within the bell during contraction required a different approach for angular measurements. Medusan angular changes were measured by changes of relatively fixed structures within the bell, the gonads, during medusan turning. The angle of the selected gonads were measured relative to the lab-fixed frame in successive images using Image J v1.48 software (National Institutes of Health).

## Supplementary Materials

**Figure S1.**
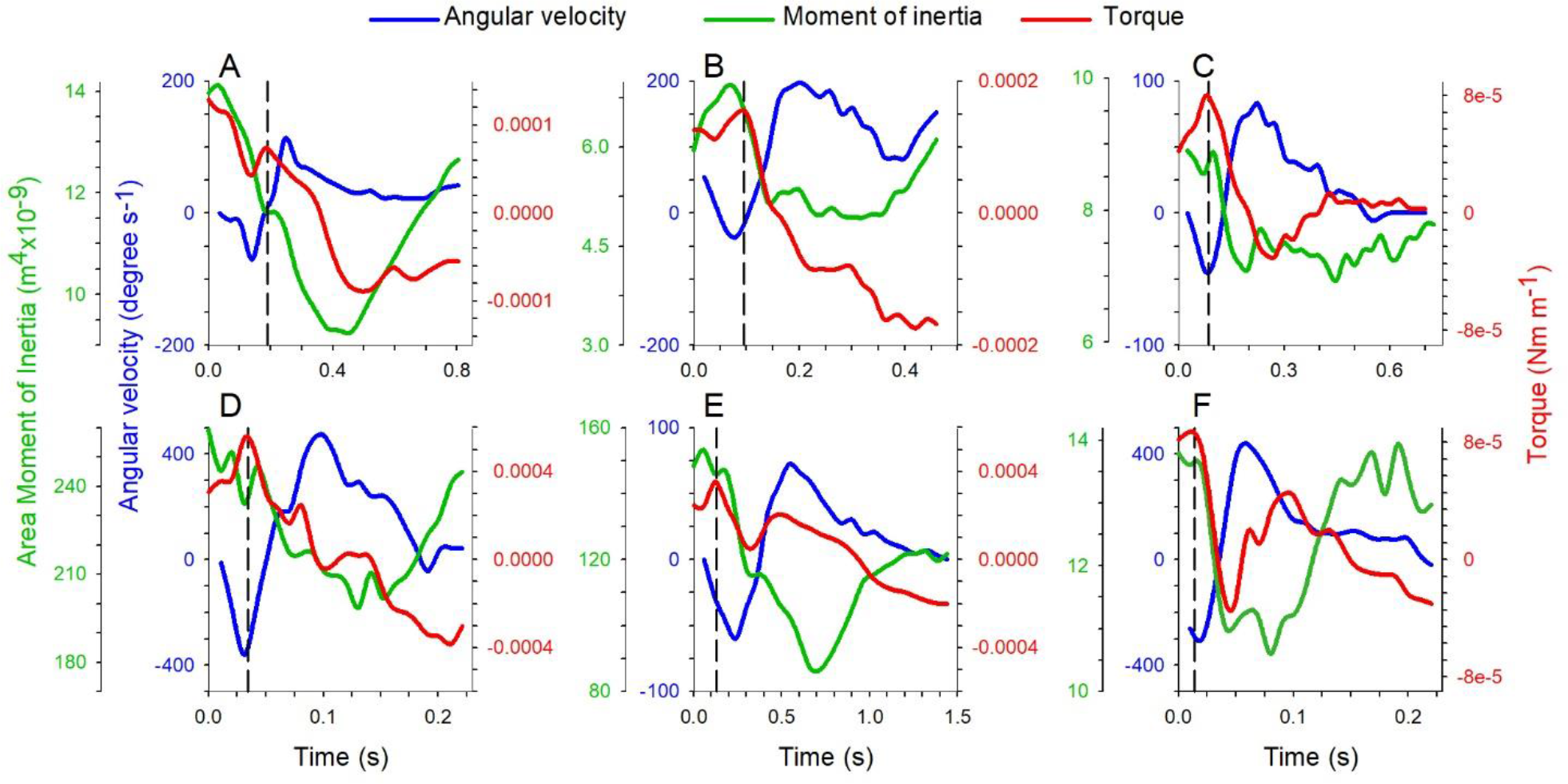
Turning parameters for medusa (*Aurelia aurita*) executing turns of different magnitude. Variable designations are same as in Fig. 1: torque per unit depth (red line), angular velocity (blue line) and moment of inertia (green line). Bell diameter and total turn angle for each turn: **A**) 2.7 cm, 53°, **B**) 1.8 cm, 50°, **C**) 2.3 cm, 13°, **D**) 4.9 cm, 30°, **E**) 5.4 cm, 20°, **F**) 2.5 cm, 23°. Local peak in torque is indicated by vertical dashed line each panel.

**Figure S2.**
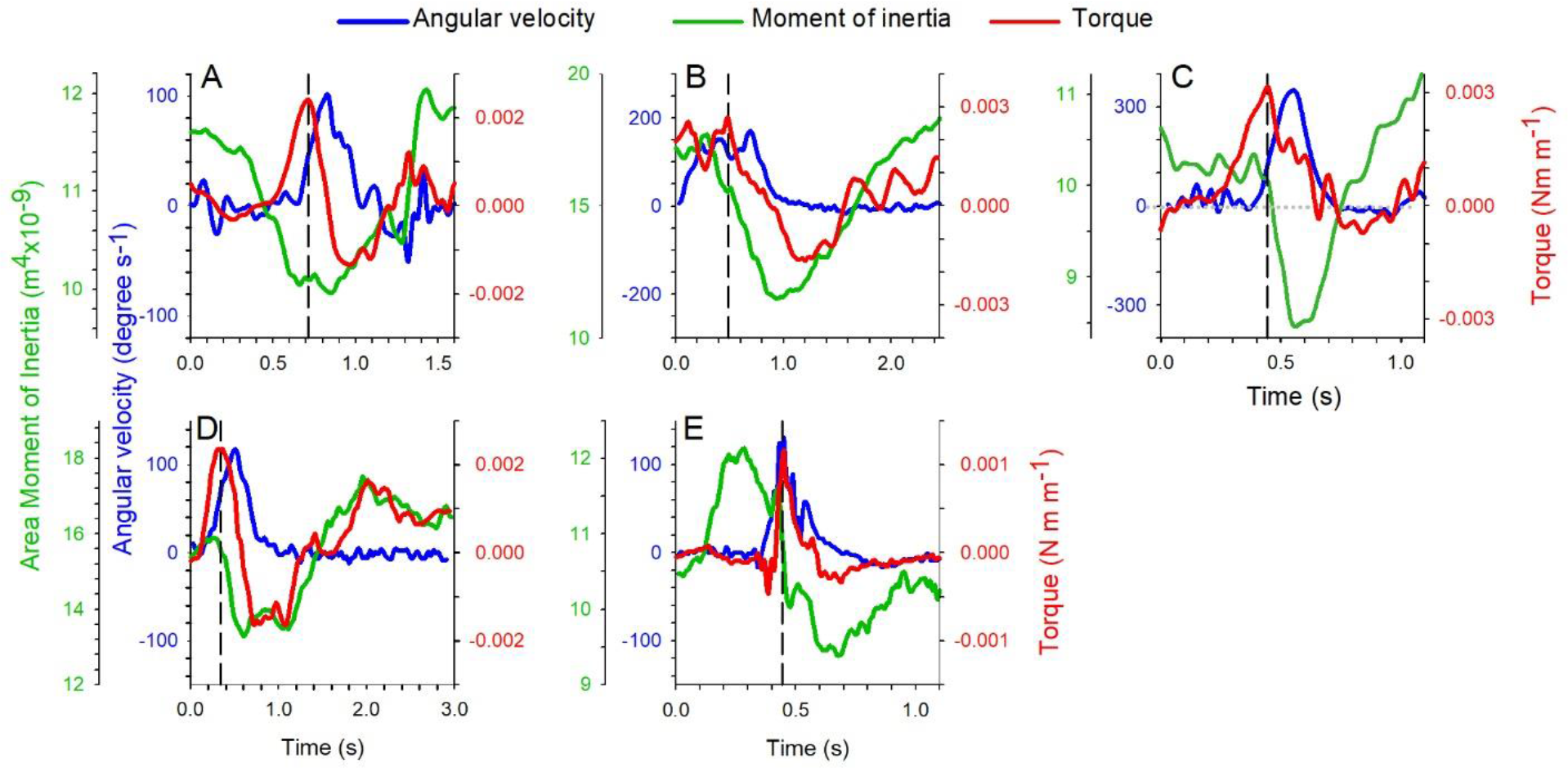
Turning parameters for zebrafish (*Danio rerio*) executing turns of different magnitude. Variable designations are same as in Fig. 1: torque per unit depth (red line), angular velocity (blue line) and moment of inertia (green line). Fish body standard length and total turn angle for each turn: **A**) 3.8 cm, 17°, **B**) 3.5 cm, 95°, **C**) 3.2 cm, 62°, **D**) 3.8 cm, 39°, **E**) 3.3 cm, 24°. Local peak in torque is indicated by vertical dashed line each panel.

Movie S1: Particle image velocimetry video sequence for turning *Aurelia aurita*.

Movie S2: Pressure video sequence for turning *Aurelia aurita*.

Movie S3: Force along the swimmer’s body video sequence for turning *Aurelia aurita*.

Movie S4: Particle image velocimetry video sequence for turning *Danio rerio*

Movie S5: Pressure video sequence for turning *Danio rerio*.

Movie S6: Force along the swimmer’s body video sequence for turning *Danio rerio*.

## Acknowledgments

We thank S. Spina and C Doller of the New England Aquarium for providing *A*. a*urelia* and J. Gitlin of the Marine Biological Laboratory for providing *Danio rerio* used in our experimental work.

## Author contributions

All authors conceived the research; SPC, BJG, MCL, and JHC collected animal measurements; all authors analyzed data; JOD and JHC wrote the initial manuscript and all authors contributed to revisions.

## Data availability

All data, including original video sequences, will be made available upon reasonable request to the corresponding author.

## References

1. J. E. Schriefer, M. E. Hale, Strikes and startles of northern pike (Esox lucius): a comparison of muscle activity and kinematics between S-start behaviors. Journal of Experimental Biology 207, 535–544 (2004).

2. P. Domenici, R. Blake, The kinematics and performance of fish fast-start swimming. Journal of Experimental Biology 200, 1165–1178 (1997).

3. P. Domenici, Context-dependent variability in the components of fish escape response: integrating locomotor performance and behavior. Journal of Experimental Zoology Part A: Ecological Genetics and Physiology 313, 59–79 (2010).

4. M. S. Triantafyllou, A. H. Techet, F. S. Hover, Review of experimental work in biomimetic foils. IEEE Journal of Oceanic Engineering 29, 585–594 (2004).

5. D. B. Quinn, G. V. Lauder, A. J. Smits, Scaling the propulsive performance of heaving flexible panels. Journal of fluid mechanics 738, 250–267 (2014).

6. S. Alben, Simulating the dynamics of flexible bodies and vortex sheets. Journal of Computational Physics 228, 2587–2603 (2009).

7. G. K. Taylor, R. L. Nudds, A. L. Thomas, Flying and swimming animals cruise at a Strouhal number tuned for high power efficiency. Nature 425, 707 (2003).

8. M. Triantafyllou, G. Triantafyllou, R. Gopalkrishnan, Wake mechanics for thrust generation in oscillating foils. Physics of Fluids A: Fluid Dynamics 3, 2835–2837 (1991).

9. C. Eloy, Optimal Strouhal number for swimming animals. Journal of Fluids and Structures 30, 205–218 (2012).

10. K. Schmidt-Nielsen, Locomotion: energy cost of swimming, flying, and running. Science 177, 222–228 (1972).

11. B. J. Gemmell et al., Passive energy recapture in jellyfish contributes to propulsive advantage over other metazoans. Proceedings of the National Academy of Sciences 110, 17904–17909 (2013).

12. M. Lighthill, Note on the swimming of slender fish. Journal of fluid Mechanics 9, 305–317 (1960).

13. S. Vogel, Life in moving fluids: the physical biology of flow. (Princeton University Press, 1996).

14. M. Sfakiotakis, D. M. Lane, J. B. C. Davies, Review of fish swimming modes for aquatic locomotion. IEEE Journal of oceanic engineering 24, 237–252 (1999).

15. N. E. Humphries et al., Environmental context explains Lévy and Brownian movement patterns of marine predators. Nature 465, 1066 (2010).

16. A. M. Reynolds, Current status and future directions of Lévy walk research. Biology open 7, bio030106 (2018).

17. G. M. Viswanathan et al., Optimizing the success of random searches. Nature 401, 911 (1999).

18. L. Seuront, F. G. Schmitt, S. Souissi, M. Brewer, J. Strickler, From random walk to multifractal random walk in zooplankton swimming behaviour. Zoological Studies 43, 498–510 (2004).

19. J. E. Stanistreet, D. Risch, S. M. Van Parijs, Passive acoustic tracking of singing humpback whales (Megaptera novaeangliae) on a Northwest Atlantic feeding ground. PLoS One 8, e61263 (2013).

20. C. Berg, J. Rayner, The moment of inertia of bird wings and the inertial power requirement for flapping flight. Journal of experimental biology 198, 1655–1664 (1995).

21. M. Thollesson, U. M. Norberg, Moments of inertia of bat wings and body. Journal of experimental Biology 158, 19–35 (1991).

22. C. Frohlich, The physics of somersaulting and twisting. Scientific American 242, 154–165 (1980).

23. P. J. Sinclair, C. A. Walker, S. Cobley, in ISBS-Conference Proceedings Archive. (2014).

24. J. H. Costello, S. P. Colin, J. O. Dabiri, Medusan morphospace: phylogenetic constraints, biomechanical solutions, and ecological consequences. Invertebrate Biology 127, 265–290 (2008).

25. K. E. Severi et al., Neural control and modulation of swimming speed in the larval zebrafish. Neuron 83, 692–707 (2014).

26. K. N. Lucas, J. O. Dabiri, G. V. Lauder, A pressure-based force and torque prediction technique for the study of fish-like swimming PLoS ONE 12, e0189225. (2017).

27. J. O. Dabiri, S. Bose, B. J. Gemmell, S. P. Colin, J. H. Costello, An algorithm to estimate unsteady and quasi-steady pressure fields from velocity field measurements. Journal of Experimental Biology, jeb. 092767 (2014).

28. T. L. Daniel, Unsteady aspects of aquatic locomotion. American Zoologist 24, 121–134 (1984).

29. D. R. Carrier, R. M. Walter, D. V. Lee, Influence of rotational inertia on turning performance of theropod dinosaurs: clues from humans with increased rotational inertia. Journal of Experimental Biology 204, 3917–3926 (2001).

30. I. Nesteruk, G. Passoni, A. Redaelli, Shape of aquatic animals and their swimming efficiency. Journal of Marine Biology 2014, (2014).

31. C. Friedman, M. Leftwich, The kinematics of the California sea lion foreflipper during forward swimming. Bioinspiration & biomimetics 9, 046010 (2014).

32. F. E. Fish et al., Kinematics of swimming of the manta ray: three-dimensional analysis of open water maneuverability. Journal of Experimental Biology, jeb. 166041 (2018).

33. X. Zhao, Designing toughness and strength for soft materials. Proceedings of the National Academy of Sciences 114, 8138–8140 (2017).

